# Cognitive reappraisal and corresponding neural basis mediate the association between childhood maltreatment and depression

**DOI:** 10.1101/2023.02.24.529872

**Authors:** Yu Mao, Ling Li, Xin Hou, Yuan Li, Shukai Duan

## Abstract

**Background:** Childhood maltreatment is considered as a robust predictor of depression. However, the underlying psychological and neurological mechanisms linking childhood maltreatment and depression remain poorly understood. Sufficient evidence demonstrates emotion dysregulation in individuals who have experienced childhood maltreatment, but it is unknown whether these changes represent vulnerability for depression. Here we speculated that decreased cognitive reappraisal and its corresponding neural basis might explain the relationship between childhood maltreatment and follow-up depression.

**Methods:** First, we investigated whether cognitive reappraisal can explain the relationship between childhood maltreatment and depression, with a cross-sectional (*n* = 657) behavioral sample. Then we recruit 38 maltreated participants and 27 controls to complete the cognitive reappraisal functional magnetic resonance imaging (fMRI) task. The between-group difference in brain activation and functional connectivity (FC) were tested using independent t-tests. Finally, we investigated the relationship between childhood maltreatment, task-based brain activity and depression.

**Results:** The behavior results suggested that cognitive reappraisal mediate the association between childhood maltreatment and depression. Specifically, participants with higher level of childhood maltreatment tend to have deficit in cognitive reappraisal, which ultimately predict higher level of depression when facing stressful life event. In addition, the maltreated group exhibited lower activation of orbitofrontal cortex (OFC) and higher FC of between the dorsolateral prefrontal cortex (DLPFC), posterior cingulate cortex (PCC), OFC, and amygdala during cognitive reappraisal, compared with healthy controls. Furthermore, the FC of DLPFC-amygdala mediates the association between childhood maltreatment and depression.

**Conclusion:** In summary, childhood maltreatment is associated with inefficient cognitive reappraisal ability, manifesting as aberrant modulation of cortical areas on amygdala. These cognitive and neural deficits might explain the relationship between childhood maltreatment and risk of depression in later life. On the other side, cognitive reappraisal might also be a potential resilient factor for the prevention of maltreatment related emotional problems.

## 1. Introduction

Previous studies had established that childhood maltreatment existed widely all over the world (Ma, 2018; Scher et al., 2004). Numerous epidemiological and clinical studies have suggested a strong relationship between childhood maltreated experiences and depression (Braithwaite et al., 2017; Nanni et al., 2012; Nelson et al., 2017). For example, individuals who experienced childhood abuse were found to have a four-fold increased risk of major depressive disorder (Felitti et al., 2019). In addition, the severity of childhood trauma is significantly associated with the duration and severity of depressive symptoms (Bifulco et al., 1991; Chapman et al., 2004; Gibb et al., 2001; Mullen et al., 1996). Maltreated children have a significantly increased risk of depression in adolescence and adulthood, reflecting the lasting and profound effects of maltreated childhood experiences on depression. In recent years, the incidence of depression has increased significantly. About one quarter of maltreated children develop severe depression during adulthood, which causes a huge health burden for society (Mello et al., 2009). This collective evidence highlights the importance of exploring early intervention in abused and neglected children before they develop depression. However, the psychological and neurological mechanisms linking childhood maltreatment and the occurrence of depression remain incompletely understood.

Individuals who experience childhood maltreatment often exhibit emotion dysregulation (Burns et al., 2010; Eisenberg et al., 2010). Childhood is considered a critical period for brain development, and maltreatment during childhood was suggested to be associated with remarkable functional and structural changes in the brain (Teicher et al., 2016). One of the most consistent findings is deficits in the limbic system, which is involved in emotion processing and regulation (Dannlowski et al., 2012). Childhood is also a dynamic period for the development of emotion regulation behavior, during which children learn to regulate emotions consciously (Bargh & Williams, 2007; Rottenberg & Gross, 2003; Silvers, 2022) in order to be able to respond effectively to environmental demands (Gross & Muñoz, 1995). However, maltreated experiences in childhood can disrupt the development of effective emotional regulation strategies, which may negatively impact subsequent emotional function (Briere et al., 2008; Shields & Cicchetti, 2001; Spasojević & Alloy, 2002). Recently, cognitive reappraisal has received increasing attention based on evidence of its effectiveness and adaptability for improved regulation of negative emotions with minimal physiological and cognitive strain (Gross & John, 2003). Indeed, habitual use of cognitive reappraisal tends to be associated with greater mental health (Gross & John, 2003) and lower levels of psychopathology (Eftekhari et al., 2009; Werner & Gross, 2010). However, study has shown that maltreated children display fewer adaptive emotion regulation strategies, such as cognitive reappraisal (Shipman et al., 2007).

In summary, the relationship between childhood maltreatment and depression has been confirmed, but few studies have focused on the psychological mechanism by which childhood maltreatment affect depression. The present study was designed to explore the psychological and neurological mechanisms underlying the link between childhood maltreatment and depression with two behavior datasets and a task-functional magnetic resonance imaging (fMRI) dataset. Such information is critical for mitigating the effects of childhood maltreatment and for developing interventions for depression. Neuroimaging evidence suggested that emotion regulation involves regulation of the amygdala by the prefrontal cortex, and the related processes include fear elimination, stress adaptation, and regulation of responses to emotional stimuli (Etkin et al., 2006; Weinberg et al., 2010). We hypothesized that deficits in cognitive reappraisal could explain the relationship between childhood maltreatment and depression. Furthermore, individuals with a history of childhood maltreatment exhibited aberrant fronto-amygdala connectivity during cognitive reappraisal, compared with controls. The observed deficit in fronto-amygdala connectivity might mediate the association between childhood and maltreatment.

## 2. Method

### 2.1. Participants

To exam the possible mediating mechanisms, we employed a cross-sectional behavioral dataset with 657 healthy college students (males: *n* = 215, age 17-25 years). Another longitudinal behavioral dataset was used to test whether cognitive reappraisal could explain the relationship between childhood maltreatment and depression under stressful life events. This dataset contained 794 college students (males: *n* = 287, age 17-26 years). They completed the measures of childhood maltreatment and cognitive reappraisal before the coronavirus disease 2019 (COVID-19) outbreak, and depression was measured after the COVID-19 outbreak (February 24, 2020).

The task-fMRI dataset contained the maltreated group and control group. Before the MRI experiment, we screened the participants using the childhood trauma questionnaire, and selected 38 participants who met the criteria as the maltreated group and 27 subjects as the control group. The screen criteria was recommended by the Childhood Trauma Questionnarie (CTQ) manual (Bernstein et al., 1997; Heim et al., 2009; Moog et al., 2018). For the maltreated group, the criteria were scores for emotional abuse > 13, physical abuse > 10, sexual abuse > 8, emotional neglect > 15, and physical neglect > 10. The control group scored no more than the standard score on each CTQ subscale. All participants were right-handed, without psychopathological symptoms or neurological diseases, and had normal or corrected visual acuity. Before the start of the experiment, the participants confirmed they understood all the experimental procedures and signed the informed consent form. To ensure the quality of the fMRI data, 8 participants were excluded for the mean framewise displacement (FD) of their data exceeded 2.5. Thus, 57 participants remained, including 33 participants in the maltreated group and 24 participants in the control group.

### 2.2. Behavior measures

#### 2.2.1. Childhood Trauma Questionnaire

The Short Form of Childhood Trauma Questionnaire (CTQ-SF) was used to assess participants’ traumatic experiences during childhood (Bernstein et al., 2003). The CTQ-SF consists of 25 questions and answers are ranked on a five-point scale (1 = never occurring to 5 = always occurring). The CTQ-SF focuses on five types of childhood trauma, including emotional, physical, and sexual abuse, and emotional and physical neglect. The CTQ-SF shows good reliability and validity (Jiang et al., 2018).

#### 2.2.2. Emotion Regulation Questionnaire

The Emotion Regulation Questionnaire is a 10-item self-report questionnaire (Gross & John, 2003) that widely used for measuring individual difference of expression suppression and cognitive reappraisal. Participants were asked to answer some questions about their emotional life, included how they control (i.e., adjustment and management) negative emotions. Previous studies have proved that the scale has high construct validity (Gross & John, 2003).

#### 2.2.3. Beck Depression Scale

The Beck Depression Inventory-II (BDI-II) was used to measure participants’ depression level. Each of the 21 items was rated on a 4-point Likert scale from 0 to 3. Participants who scored higher on the BDI showed more depressive symptoms (Beck et al., 1996). The Baker Depression Scale includes cognitive emotional factors and somatic factors. Cognitive affective factors include sadness, experience of failure, loss of pleasure, guilt, punishment, self-loathing, self-criticism, suicidal thoughts, crying, emotional arousal, loss of interest, indecision, feelings of worthlessness, and irritability. Somatic factors include energy loss, changes in sleep patterns, changes in appetite, concentration, and tiredness or fatigue (Beck et al., 1996; Dozois et al., 1998). The BDI-II is a reliable and widely used measure to assess the severity of depressive symptoms in non-clinical and clinical samples.

#### 2.2.4. Emotion regulation paradigm

All participants completed the emotion regulation task, which was adapted from a previously validated paradigm (Minkel et al., 2012; Ochsner et al., 2004) under MRI observation. This task contained two emotion regulation strategies: cognitive reappraisal and expressive suppression. Cognitive reappraisal refers to the alteration of negative emotions by changing the way one thinks, while expressive suppression indicated suppressing the expression of emotions. The task was constructed with four conditions, including viewing-natural image (15 trials), viewing-negative image (15 trials), reappraisal-negative image (15 trials) and suppression-negative image (15 trials). The image were selected from the International Affective Picture System (IAPS) database (The International Affective Picture System (IAPS) for the study of emotion and attention;. The valence, arousal and dominance values of negative images were matched across each condition.

### 2.3. MRI data acquisition and preprocessing

The whole brain images of the participants were acquired by a Siemens 3.0T scanner (Siemens Medical Systems, Erlangen, Germany) using a 32-chanel brain coil at Southwest University Brain Imaging Center. Functional images were obtained using EPI sequence: TR = 2,000 ms, echo TE = 30 ms, FOV = 224 × 224, FA = 90°, slices = 32, thickness = 3 mm, slice gap = 1 mm, voxel size = 3.4 × 3.4 × 3 mm^3^.

Neuroimaging data were preprocessed using Statistical Parametric Mapping (SPM8, http://www.fil.ion.ucl.ac.uk/spm). The first 4 volumes were discarded to suppress the equilibration effects and consider subjects’ adaptation to the environment. The remaining volumes were slice-timing corrected, realigned, and spatially normalized to the MNI (Montreal Neurological Institute) template and resampled to 2 × 2 × 2 mm^3^. We next conducted spatial smoothing with 6-mm full-width at half-maximum Gaussian kernel.

### 2.4. GLM model

Then we constructed a general linear model for each participant with canonical hemodynamic response functions. The present study focused on the neural activity underlying cognitive reappraisal. Each cognitive reappraisal specific brain map was obtained by contrasting the cognitive reappraisal condition with viewing negative image condition. In second-order analyses, we focused on the differences in brain activation between the maltreated and control groups under cognitive reappraisal condition using independent sample t-tests. Significant brain regions were identified based on Family-Wise error (FWE) correction of the nonstationary superthreshold cluster size distribution calculated by Monte Carlo simulation with an intensity threshold of P <0.001 and a breadth threshold of P <0.05 (Nichols & Hayasaka, 2003).

### 2.5. PPI analysis

Psychophysiological interaction (PPI) analyses were used to explore the group difference in task-induced connectivity between brain regions during cognitive reappraisal. Considering the critical role of the amygdala in emotion processing, the bilateral amygdala was chosen as the region of interest (ROI) for the PPI analyses. In addition, the significant clusters of the group comparison in activation analyses were chosen as ROI. We then used independent sample t-tests to investigate the group difference in task-based functional connectivity (FC). The results were corrected by FWE correction of nonstationary superthreshold cluster size distribution calculated by Monte Carlo simulation with an intensity threshold of P <0.001 and a breadth threshold of P <0.05 (Nichols & Hayasaka, 2003).

### 2.6. Mediation analyses

To test whether cognitive reappraisal and its corresponding neural basis could explain the relationship between childhood maltreatment and depression, we performed a mediation analysis using the indirect macro designed for SPSS (Preacher & Hayes, 2008). The mediation analyses were employed in two behavior sample separately. X is the childhood maltreatment, Y is the depression, and M is cognitive reappraisal and its corresponding neural basis. Age and sex were used as covariates in the model. This macro uses bootstrapped sampling to estimate the indirect mediation effect. In this analysis, 2000 bootstrapped samples were drawn, and bias corrected 95 % bootstrap confidence intervals (CIs) were reported. CIs that do not include zero indicate a significant indirect effect of the independent variable on the dependent variable through the mediators (Preacher & Hayes, 2008).

### 2.7. Prediction analyses

To test the robustness of the brain-behavior relationship, we employed a machine-learning method called linear support vector regression (SVR) with leave one out cross-validation procedure (Supekar et al., 2013). Depression was taken as the dependent variable and the amygdala-seeded FC during cognitive reappraisal as independent variables in the linear regression algorithm. The r_(predicted, observed)_ was estimated by leave one out cross-validation, and represented the predictive accuracy of the independent variable.

## 3. Results

### 3.1. Sample characteristics and behavior results

In the cross-sectional behavioral sample, there are significant correlations between childhood maltreatment, depression and cognitive reappraisal (Table 1). The mediating model of emotion regulation showed that cognitive reappraisal could mediate the relationship between childhood maltreatment and depression (indirect effect = 0.02, 95% CI = [0.037,0.417], *P* = 0.03, Figure 1a). In the longitudinal sample, significant correlations were observed between childhood maltreatment, depression and cognitive reappraisal (Table 1). The mediating model also revealed that cognitive reappraisal mediated the relationship between childhood maltreatment and depression under stressful life event (indirect effect = 0.02, 95% CI = [0.015, 0.052], *P* < 0.001, Figure 1b), suggesting the potential of cognitive reappraisal as an intervening mechanism to attenuate the tight relationship between childhood maltreatment and depression.

**Table 1.** The correlation coefficients between childhood maltreatment, depression and cognitive reappraisal.

| Sample | Variable | Childhood maltreatment | Cognitive reappraisal | Depression_T1 | Depression_T2 |
| --- | --- | --- | --- | --- | --- |
| Cross-sectional sample | Childhood maltreatment | - | -0.126*** | 0.203*** | - |
|  | Cognitive reappraisal | - | - | --0.129*** | - |
| Longitudinal sample | Childhood maltreatment | - | -0.170*** | 0.339*** | 0.296*** |
|  | Cognitive reappraisal | - | - | -0.303*** | -0.255*** |

\*\*\* $P < 0.001$

**Figure 1.**
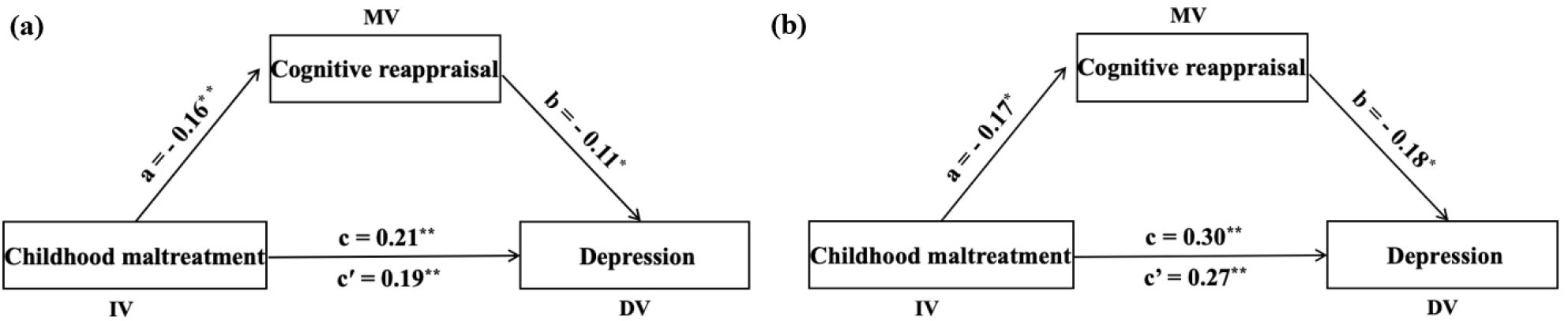
Cognitive reappraisal mediates the relation between childhood maltreatment and depression. (a) represents the result from cross-sectional sample and (b) represents the result from the longitudinal sample.

Independent sample t-tests showed significant differences in childhood trauma scores (t = −7.338, p = 0.001) and depression scores (t = −5.090, *P* = 0.002) between the maltreated group and control group (Table 2). No significant difference in age was observed between the two groups (t = 0.235, *P* = 0.520). The self-rating negative emotion in cognitive reappraisal condition and viewing negative image condition were 2.423 and 3.611, respectively. Paired sample t-test showed that the self-rating negative emotion score in the condition of viewing negative images was significantly higher than that in the condition of cognitive reappraisal (t =12.148, *P* < 0.001), indicating that the participants successfully reduced negative emotions by cognitive reappraisal. The correlation analysis showed that childhood maltreatment was significantly correlated with depression (r = 0.490, *P* = 0.01) and cognitive reappraisal (r = − 0.307, *P* = 0.02), and cognitive reappraisal was significantly correlated with depression (r = − 0.371, *P* = 0.003). Further mediating analysis showed that cognitive reappraisal mediated the relationship between childhood maltreatment and depression (95% [CI: 0.067,0.123]), as shown in Figure 3a.

**Table 2.** The demographic data of maltreated group and control group.

| | CT group ($n = 38$) | Control group ($n = 27$) | $T$ value | $P$ value |
| --- | --- | --- | --- | --- |
| age | 19.23±0.91 | 19.51±0.72 | -0.30 | 0.371 |
| CT score | 40.09±6.07 | 32.15±4.28 | -6.77 | 0.021 |
| Depression score | 8.32±5.68 | 7.82±7.72 | -0.24 | 0.002 |

**Figure 2.**
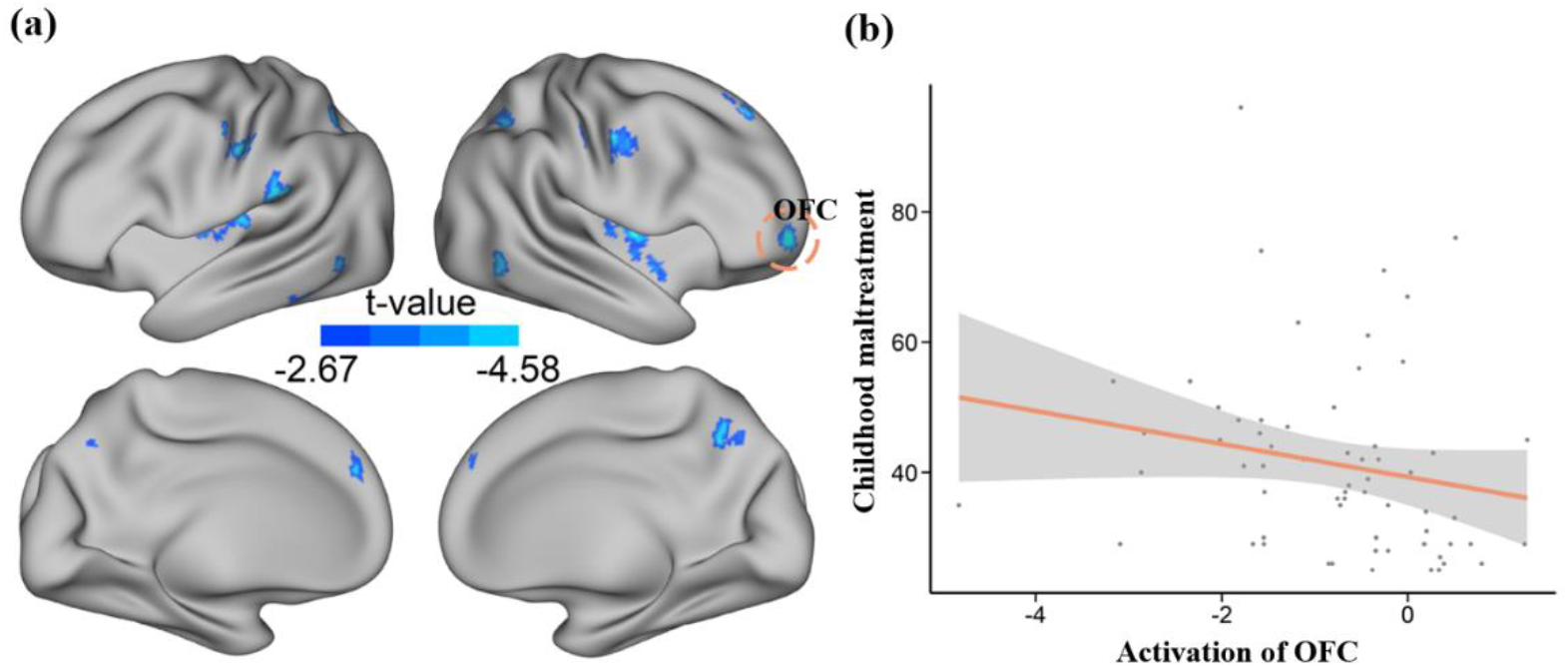
Intergroup differences of brain activation. (a) Maltreated group demonstrated lower OFC activation during cognitive reappraisal, compared with control group. (b) The activation of OFC was significantly correlated with childhood maltreatment.

**Figure 3.**
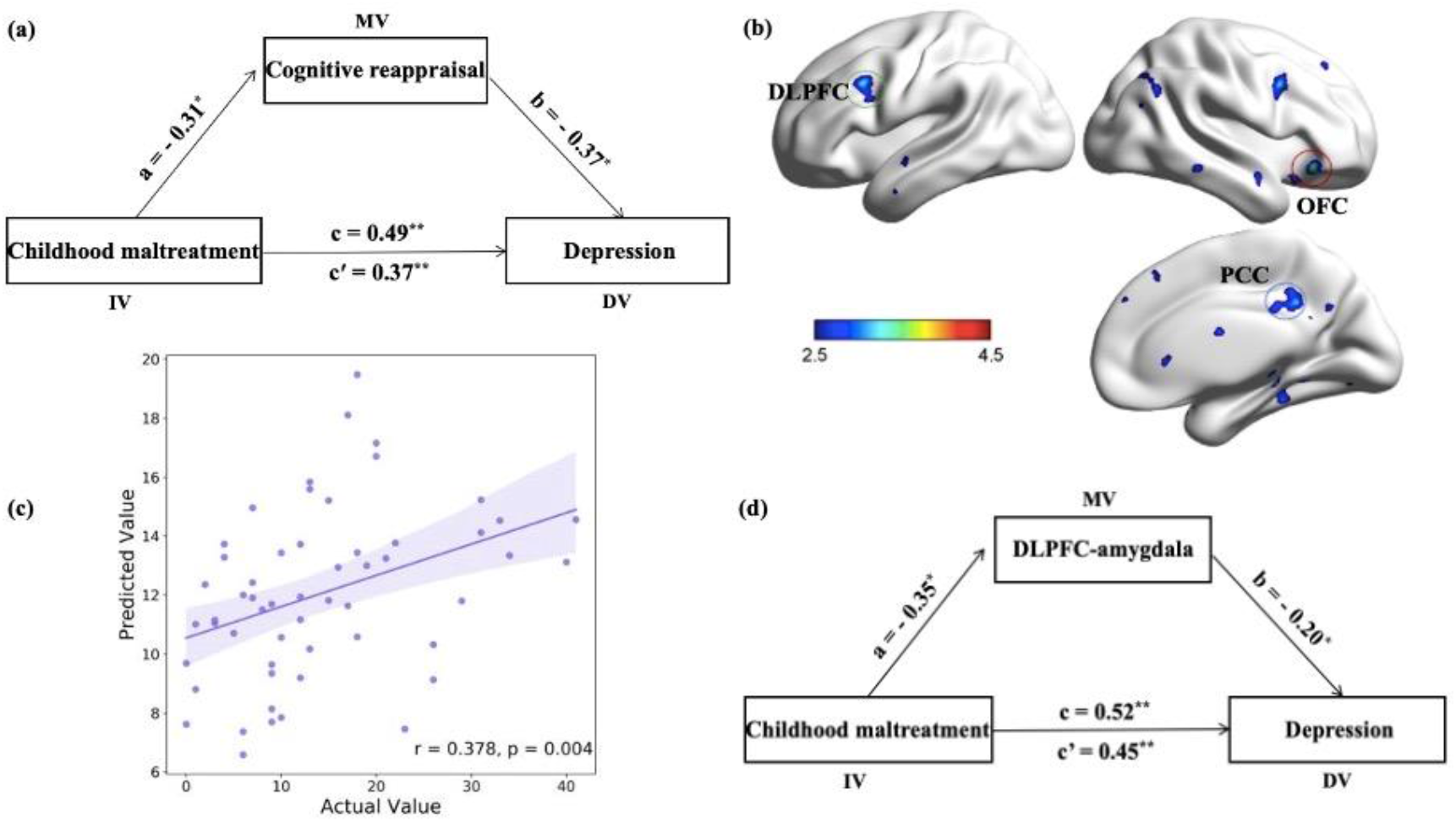
The relationship between childhood maltreatment, amygdala-seeded FC and depression. (a) Cognitive reappraisal mediates the relationship between childhood maltreatment and depression in the fMRI task. (b) Maltreated group demonstrated higher FC between left amygdala and left DLPFC (green circle), left DLPFC (green circle), left DLPFC (green circle) in cognitive reappraisal, compared with control group. (c) The FC of DLPFC-amygdala, PCC-amygdala could predict participants’ depression. (d) The FC between DLFPC and amygdala mediates the relationship between childhood maltreatment and depression.

### 3.2. Activation results

We next investigated the differences in brain activation patterns between the maltreated group and the control group in the cognitive reappraisal condition. The results revealed significant differences in the activation intensity in orbitofrontal cortex (OFC, x = 28, y = 54, z = −4, T = −4.58) between the two groups (Figure 2). Then we extracted the activation intensity value in the OFC and analyzed the relationship between the activation intensity in the OFC, childhood maltreatment and depression.

### 3.3. PPI results

PPI analysis showed that in the cognitive reappraisal condition, maltreated group and control group exhibited significant differences in the FC of the left dorsolateral prefrontal cortex (DLPFC) - left amygdala (x = −46, y =18, z = 34, T = 4.44), the right posterior cingulate cortex (PCC) - left amygdala (x = 2, y =- 40, z =34, T=3.71) and the right OFC - left amygdala (x = 42, y =30, z = −12, T = 3.78) (see Figure 3a).

### 3.4. Brain-behavior associations

Pearson correlation analyses showed that the activation intensity in the OFC was significantly correlated with childhood maltreatment (r = −0.301, *P* = 0.030). Moreover, there significant associations between childhood maltreatment and left DLPFC - left amygdala (r = 0.356, *P* =0.010), the right PCC - left amygdala (r = 0.396, *P* = 0.004) and the right OFC - left amygdala (r = 0.358, *P* =0.009). Depression was significantly correlated with the FC of left DLPFC - left amygdala (r = 0.350, *P* =0.010), the right PCC - left amygdala (r = 0.406, *P* =0.003, Figure 3b). The prediction model based on machine learning showed that the FC between the right PCC and left amygdala and between the left DLPFC and left amygdala during cognitive reappraisal was significantly correlated with depression (r=0.378, *P* =0.004, as shown in Figure 3c). Moreover, the FC of DLPFC-amygdala mediate the association between childhood maltreatment and depression (95% [CI: 0.203, 0.163]), as shown in Figure 3d.

## 4. Discussion

The present study investigated the psychological and neurological mechanisms underlying the association between childhood maltreatment and depression. First, we confirmed the mediating role of cognitive reappraisal linking childhood maltreatment and depression with a cross-sectional and a longitudinal sample. Then, an emotional regulation task was used to investigate the differences in brain activation patterns between the maltreated group and the control group during cognitive reappraisal. The results demonstrated significant between-group differences in activation of the OFC and in the FC between the DLPFC, OFC, PCC and amygdala during cognitive reappraisal. Subsequent correlation analysis revealed that activation of the OFC, as well as the FC of the DLPFC, PCC and amygdala were significantly correlated with childhood maltreatment. Further predictive analysis revealed that the FC of the DLPFC and OFC to the amygdala were significantly associated with depression. In summary, the integrated evidence obtained in this study through both behavioral and neuroimaging results suggests a mediating role of cognitive reappraisal in the link between childhood maltreatment and depression.

### 4.1. The association between childhood maltreatment, cognitive reappraisal and depression

Cognitive reappraisal is suggested to be one of the most adaptive emotion regulation strategy, and can effectively reduce depressive symptoms and improve life satisfaction (Garnefski et al., 2004; Garnefski & Kraaij, 2006; Kashdan & Steger, 2006). Empirical studies suggested that using cognitive reappraisal to regulate negative emotion would not impair memory or increase physiological arousal, such as blood pressure or heart rate (Egloff et al., 2006; Richards & Gross, 2000), especially, an experimental study demonstrated that the application of cognitive reappraisal strategy during emotion regulation can reduce the physiological arousal of individuals (Dandoy & Goldstein, 1990). The positive effect of cognitive reappraisal has been investigated in clinical and preclinical populations, and the results demonstrated that the use of cognitive reappraisal strategies is relatively higher in healthy subjects than in patients with mental disorders (Bryant et al., 2001; Garnefski et al., 2002). The present study revealed that cognitive reappraisal mediated the association between childhood maltreatment and depression, which in turn imply that cognitive reappraisal may be a modifiable target for the intervention of childhood maltreatment-related emotional problems.

### 4.2. The association between childhood maltreatment, cognitive reappraisal corresponding neural basis and depression

The maltreated group showed lower OFC activation during cognitive reappraisal, compared with control group. In addition, the activation intensity in the OFC was negatively correlated with the childhood trauma scores. Previous studies have confirmed the role of the OFC in emotion evaluation and regulation (Golkar et al., 2012; Rudebeck & Murray, 2014; Wager et al., 2008). The OFC receives and monitors neural inputs from a variety of sensory modalities, including olfactory information (Zatorre et al., 1992), auditory information (Frey et al., 2000) and somatic sensation (Small et al., 2007). In addition, the OFC can directly project to the primary and secondary sensory cortex, which facilitates the modulation of neural signals from the sensory cortex and ultimately influences the individual’s behavior (Kringelbach & Rolls, 2004). Previous research also suggested that the neural connections between the OFC and subcortical structures (amygdala, thalamus, etc) are crucial pathways underlying emotion processing (Wager et al., 2008). Specifically, a study using fMRI task analysis showed that the presence of interactions between specific regions of the frontal cortex (dorsolateral prefrontal, dorsomedial prefrontal, anterior cingulate, orbitofrontal) and the amygdala was observed during cognitive reappraisal (Banks et al., 2007), and the strength of amygdala connectivity to the OFC and medial prefrontal cortex could predict the reduction of negative effects after reassessment. Another study also observed FC between the OFC and amygdala during cognitive reappraisal, and increased OFC activation was associated with decreased amygdala activity (Drabant et al., 2009). In the present study, the reduced OFC activation in the maltreated group during the process of cognitive reappraisal may indicate that these individuals cannot employ cognitive reappraisal effectively through the regulation of the limbic system, when facing negative life events.

The DLPFC is a core region involved in executive control (Miller & Cohen, 2001) and the autonomous regulation of emotions (Ochsner & Gross, 2005; Phillips & Dudík, 2008). FC of DLPFC-amygdala was suggested to be a fundamental pathway underlying emotion regulation (Ochsner et al., 2012). Clinical research shown that the FC between the DLPFC and amygdala is significantly increased in depressed individuals compared with that in healthy individuals (Jin et al., 2011). The current literature suggests that childhood maltreatment is associated with enhanced prefrontal amygdala coupling in adolescents, which may be the underlying neural mechanism of individual adaptation to maltreatment (Herringa et al., 2016). As the central region of the default mode network, the PCC was reported to be involved in emotional-related processing, including emotional evaluation (Maddock et al., 2003) and regulation, via a modulatory influence on amygdala (Hahn et al., 2011; Pessoa et al., 2005). Patients with major depression disorder demonstrated abnormally elevated PCC-amygdala FC, which was suggested to be the trait vulnerability for depression (Fang et al., 2013). Collectively, these findings suggest that increased FC of the fronto-limbic circuits in response to negative stimulus may be an important mechanism of emotional adaptation after exposure to childhood maltreatment. However, although these changes may help individuals adapt to the maltreatment condition, they also may hinder the long-term development of emotional regulation.

### 4.3. Conclusion

In conclusion, the results of the present study identified cognitive reappraisal and altered connectivity within the fronto-limbic circuit as underlying factors in the relationship between childhood maltreatment and depression. Participants who experienced maltreatment during childhood exhibited significant deficits in cognitive reappraisal, including even those without a diagnosed mental disorder. Our findings also suggest that behavioral training and neural modulation focusing on cognitive reappraisal and its neural markers might provide effective intervention for childhood maltreatment related mental disorders.

authorship contribution statement

